# Extent of EMT promotes invasive, contact-induced sliding on progressively narrower fiber-like tracks

**DOI:** 10.1101/044800

**Authors:** Daniel F. Milano, Senthil K. Muthuswamy, Anand R. Asthagiri

## Abstract

Epithelial-mesenchymal transition (EMT) is a complex process by which cells acquire invasive properties that enable escape from the primary tumor. Complete EMT, however, is not required for metastasis: circulating tumor cells exhibit hybrid epithelial-mesenchymal states, and genetic perturbations promoting partial EMT induce metastasis *in vivo*. An open question is whether and to what extent intermediate stages of EMT promote invasiveness. Here, we investigate this question, building on recent observation of a new invasive property. Migrating cancer cell lines and cells transduced with prometastatic genes slide around other cells on spatially-confined, fiber-like micropatterns. We show here that low-dosage/short-duration exposure to TGFβ induces partial EMT and enables sliding on narrower (26 µm) micropatterns than untreated counterparts (41 µm). High-dosage/long-duration exposure induces more complete EMT, including disrupted cell-cell contacts and reduced E-cadherin expression, and promotes sliding on the narrowest (15 µm) micropatterns. These results demonstrate that EMT is a potent inducer of cell sliding, even under significant spatial constraints, and EMT-mediated invasive sliding is progressive, with partial EMT promoting intermediate sliding behavior. Our findings suggest a model in which fiber maturation and partial EMT work synergistically to promote invasiveness during cancer progression.

## Introduction

The epithelial-mesenchymal transition (EMT) plays a significant role in cancer progression and is often associated with metastasis and poor prognosis(1-5). EMT is a complex process in which epithelial cells lose expression of epithelial genes, upregulate mesenchymal markers and proteins associated with migration and invasion, and undergo changes in morphology (6-8). It is increasingly evident that EMT can also progress only partially, resulting in hybrid epithelial/mesenchymal (E/M) states in which cells exhibit a mix of epithelial and mesenchymal molecular markers (9-11). Indeed, complete EMT is not absolutely essential for metastasis, as evidenced for example by circulating tumor cells (CTCs) exhibiting partial EMT (12-14).

Hybrid E/M states also occur during the temporal progression of EMT induced by cues, such as transforming growth factor beta (TGFβ). TGFβ is a potent inducer of EMT(15) and promotes gradual changes in cell morphology and gene expression over the course of several days in culture. During this timespan, cells exhibit a mix of epithelial and mesenchymal molecular markers(16). After 48 hours of TGFβ treatment, non-transformed mammary epithelial MCF-10A cells upregulate smooth muscle actin. However, an overt EMT, defined by loss of E-cadherin expression, induction of vimentin expression, and a distinct morphology change, required 6 days of sustained exposure to TGFβ (16).

Given the prevalence of partial EMT and its significant role in cancer progression, it is important to better understand how cell behaviors associated with metastasis and invasion change as cells progress through the intermediate stages of EMT. As cells switch their molecular expression profile, are there gradual concomitant changes in invasive cell behavior? Do intermediate stages of EMT correspond to intermediate levels of invasive cell behavior that, despite being sub-maximal, may be quantitatively sufficient for disease progression?

To begin to examine the quantitative relationship between degree of EMT and invasive cell behavior, we investigate here the effect of TGFβ-induced EMT on a cell migration phenotype that is relevant to invasion in a fibrillar, crowded tumor microenvironment and that we recently demonstrated (17) to be correlated with metastasis. Cancer cells invade along fibers in a highly dynamic microenvironment with a high concentration of cells(18-20). How invading cancer cells respond to encounters with other cells along their migratory path will influence the efficiency of cancer cell dispersion. A cell that halts and reverses direction at every cell-cell encounter has a lower rate of dispersion compared to a cell that circumnavigates or slides around other cells to maintain its direction of movement.

We recently showed that metastatic MDA-MB-231 and BT-549 cells are highly effective in sliding around cell-cell encounters on fiber-like adhesive tracks, even on tracks much narrower than the cell diameter(17). In contrast, non-transformed mammary epithelial MCF-10A cells largely reverse direction upon encountering another cell. Corroborating the relationship between contact-induced sliding and invasive potential, molecular perturbations associated with cancer progression, including treatment with TGFβ, downregulation of E-cadherin and PARD3 tumor suppressors, and ErbB2 induction, enhanced sliding behavior in MCF-10A cells.

To what extent EMT is involved in promoting sliding behavior is unclear. Extended treatment with a high dose of TGFβ promotes EMT and enhances sliding behavior of MCF-10A cells(17). On the other hand, without inducing overt EMT, downregulation of PARD3 alongside induction of ErbB2 signaling enhanced sliding and induced metastasis in mouse models. These observations suggest that complete EMT is not required for sliding and pose the hypothesis that the degree of EMT quantitatively affects the extent of invasive sliding behavior.

Here, we sought to test this hypothesis and investigate the quantitative relationship between EMT and contact-induced sliding. We used different doses and durations of TGFβ treatment to modulate the extent of EMT and measured the corresponding ability of cells to undergo slide responses on spatially-confined, fiber-like micropatterns. Our results show that the acquisition of invasive sliding behavior is progressive and correlates with the extent of TGFβ-induced EMT. These results suggest a model wherein the extent of EMT cooperates with fiber maturation to promote an invasive phenotype during cancer progression.

## Materials and Methods

### Micropatterning and surface preparation

Fiber-like micropatterned surfaces were prepared and seeded as previously described(17). Briefly, fibronectin was adsorbed on poly(dimethylsiloxane) (PDMS) elastomeric stamps containing negative relief features and then transferred to a plasma-treated PDMS spin-coated glass surfaces with high spatial control. Patterned surfaces were visualized via fluorescence microscopy by copatterning trace Alex Fluor 594-conjugated BSA (Invitrogen, Carlsbad CA). Prior to cell seeding, surfaces were incubated with Pluronic F127 (EMD Biosciences, San Diego CA) to prevent non-specific cell adhesion. TGFβ-treated and untreated MCF-10A cells were seeded on the surface at a density of 2.0 x 10^4^ cells/mL and incubated for approximately 1 hour. Non-adhered cells were gently removed via consecutive PBS washes before the remaining adhered cells in the dish were given fresh growth media and the dish was transferred to the microscope for imaging.

### Image acquisition and analysis

Dishes were secured on a heated microscope stage maintained at 37C and 5% CO2. Individual positions were specified as x,y,z coordinates in Axiovision or Zen software (Carl Zeiss, Göttingen, Germany). Phase contrast images of every position were acquired at 5-minute intervals for 21 h at 10x magnification on an Axiovert 200M or LSM 700 confocal microscope (Carl Zeiss, Göttingen, Germany). Frame-by-frame image analysis was performed in Fiji (National Institutes of Health, open-access).

### Cell Culture

Non-transformed mammary epithelial MCF-10A cells were transfected with a double reporter to express GFP-tagged H2B and mCherry-tagged GM130 proteins as previously described(21). These 10A cells were maintained using standard 10A cell culture methods, as previously described (22). To induce EMT in these 10A cells, growth medium was supplemented with transforming growth factor-beta 1 (TGFβ, Peprotech). Two doses of TGFβ were analyzed: 5 and 20 ng/mL. The duration of TGFβ treatment ranged from 3-12 days. During extended treatments (>3 days), treated cells were passaged regularly and provided with fresh TGFβ-supplemented growth medium.

### Western Blot

E-cadherin was probed using a 1:1,000 dilution of primary antibodies (Santa Cruz Biotechnology) and a 1:1500 dilution of HRP-conjugated secondary antibody (Fisher Scientific). Erk2 was probed using a 1:200 dilution of primary antibody (Santa Cruz Biotechnology) and a 1:100,000 dilution of HRP-conjugated secondary antibody (Fisher Scientific). Western blots were imaged by ECL (Thermo Scientific) using a ChemiDoc Touch Imaging System (BioRad). Quantification of the blot by densitometry was performed in Image Lab (BioRad).

### Data analysis and statistics

Data analysis was conducted in MATLAB (Mathworks, Natick MA).

## Results and Discussion

### TGFβ-induced morphology changes are sensitive to dose and duration of treatment

Non-transformed mammary epithelial MCF-10A cells were exposed to either a low (5 ng/mL) or high (20 ng/mL) dose of transforming growth factor-beta 1 (Peprotech) for up to 12 days. Cells were passaged at 3-day intervals as previously described(22) and provided with fresh TGFβ.

Cells exposed to the high dose of TGFβ undergo a morphology change prior to 3 days of treatment (Fig. 1I, asterisks). Meanwhile, cells treated with the low 5 ng/mL dose required 6 days to demonstrate the same elongated, spindle-shape morphology consistent with EMT (Fig. 1F, asterisks). Cell-cell contacts in the high 20 ng/mL dose cohort were poorly defined by day 6 and persisted through day 12 (Fig. 1L, arrowheads). Although cells treated with the low dose undergo a morphology change, contacts between these cells remained intact throughout the entire 12-day duration of treatment (Fig. 1G-H). Control cells that were not exposed to TGFβ maintained the cobblestone morphology characteristic of epithelial cells (Fig. 1A-D). Taken together, these findings confirm that TGFβ induces a gradual progression from an epithelial to mesenchymal morphology in a dosage- and time-dependent manner.

**Figure 1.**
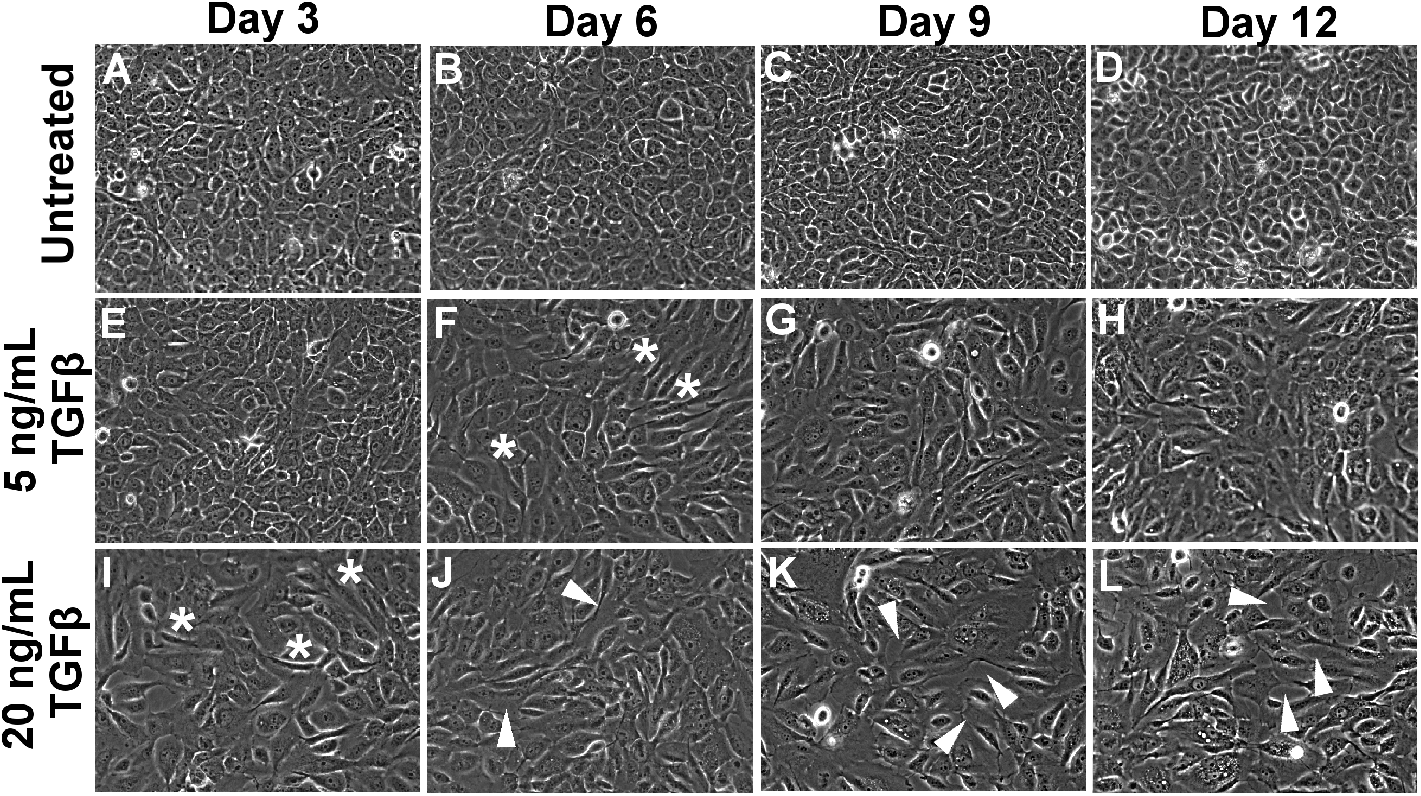
Cell morphology changes induced by TGFβ treatment. Non-transformed mammary epithelial MCF-10A cells were treated with TGFβ or left untreated. Treated cells were exposed to either 5 or 20 ng/mL of TGFβ for between 3 and 12 days. Asterisks denote cells with an elongated, mesenchymal morphology. Arrowheads indicate deficient cell-cell adhesions. Phase contrast images were taken prior to cell passage at 10x magnification.

### TGFβ-treated MCF-10A cells slide more efficiently than untreated cells

We next asked how different stages of progression along the TGFβ-induced EMT pathway affects invasive cell sliding behavior. Pairwise interactions between migrating cells with and without TGFβ treatment were imaged and the extent to which interacting cells reversed direction versus slide past each other was quantified as described previously(17).

Untreated cells are unable to slide effectively on narrow and intermediate line widths (6-21 µm) (Fig. 2A). On broader line widths, untreated cells begin to exhibit a modest level of sliding, with sliding observed in 20% of collisions at the broadest 33 µm micropattern width. These results are consistent with our previous findings showing a limited ability of non-transformed cells to undergo invasive contact-initiated slide responses(17).

**Figure 2:**
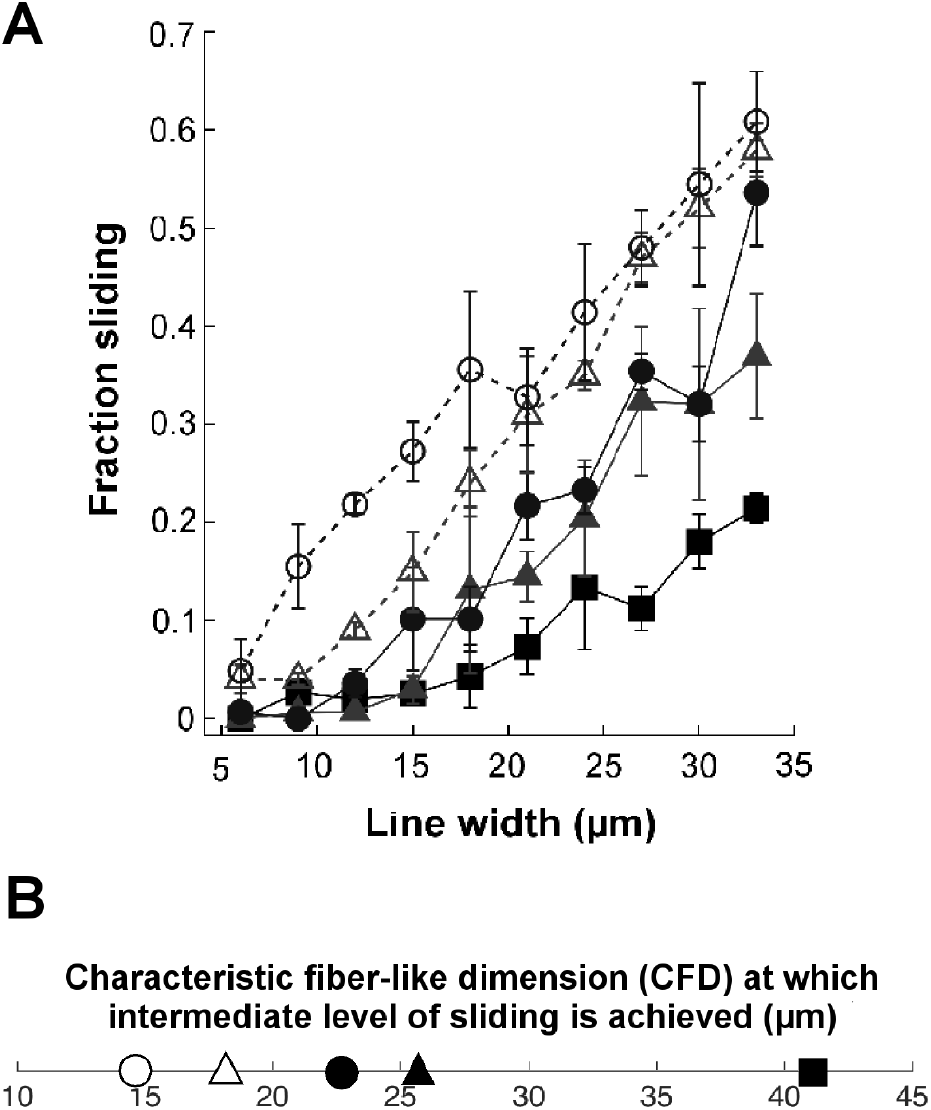
Collision response as a function of TGFβ-treatment. (*A*) Fraction sliding and (*B*) Characteristic fiber-like dimension (CFD) for TGFβ-treated and untreated (control, square) cells. MCF-10A cells were treated with either 5 ng/mL TGFβ (triangle) for 3 days (filled, solid line) or 6 days (open, dashed line), 20 ng/mL TGFβ (circle) for 3 days (filled, solid line) or 12 days (open, dashed line). Error bars denote S.E.M.

Treatment with TGFβ enhances contact-initiated sliding in a manner that depends on the dose of TGFβ, the duration of TGFβ exposure, and the micropattern width. At the higher range of micropattern widths (24-33 µm), even the shortest exposure (3 day) and lowest dose (5 ng/ml) of TGFβ increases the fraction of collisions that result in sliding. Treatment with either 5 ng/mL or 20 ng/mL TGFβ for 3 days generated similar levels of sliding. Meanwhile, increasing exposure time to 6 and 12 days for 5 ng/mL and 20 ng/mL doses of TGFβ, respectively, further increased the level of sliding. Thus, the duration of exposure has a larger effect than TGFβ dose on wide micropatterns. At the lower line widths (6-18 µm), short exposures of 3 days to TGFβ only modestly enhanced sliding behavior, if at all. Extended duration of exposure (6-12 d) was needed to substantially increase sliding behavior.

Notably, given a high TGFβ dose (20 ng/mL) and extended exposure (12 days), sliding was observed in nearly 20% of collisions on narrow 9 µm line widths. For comparison, this level of sliding was observed in untreated cells only when micropatterns were widened to 30+ µm.

To better quantify how TGFβ-induced EMT enhances cell sliding under spatial constraints, we determined the characteristic fiber-like dimension (CFD) at which cells exhibit an intermediate level of sliding. We previously demonstrated that the CFD is an effective metric for quantifying the effects of molecular perturbations on the sliding ability of non-transformed and metastatic breast epithelial cells(17). Cells that slide poorly have a high CFD and require much wider fiber-like tracks to execute moderate levels of sliding; in contrast, cells that slide proficiently have a low CFD and achieve moderate levels of sliding at low fiber-like dimensions. Since the fraction of collisions that exhibit a sliding response varied between 0% to a maximum of approximately 50-60% (Fig. 2A), we set the intermediate level of sliding to be 25%. Using a linear fit, we determine the CFD at which 25% of collisions result in a sliding response (Supplemental Fig. S1).

Untreated (control) cells have a CFD of 41 µm (Fig. 2B). Cells exposed to 5 ng/mL of TGFβ for 3 days have a lower CFD of 26 µm, indicating an enhanced ability to slide in confined environments. Extending 5 ng/mL TGFβ exposure to 6 days further reduces the CFD to 18 µm, corresponding with the acquisition of a more elongated morphology (Fig. 1). This progressive reduction in CFD with further progression along the EMT pathway is also observed at the higher TGFβ dose. At 20 ng/mL TGFβ for 3 days, cells exhibit an elongated morphology and the CFD of 23 µm is lower than the untreated control. Extending the exposure to 12 days when cell-cell contacts are compromised (Fig. 1, white arrowheads), the CFD drops even further to 15 µm.

### TGFβ-induced changes in E-cadherin expression tune collision response behavior

We next investigated the relationship between the progressive enhancement in cell sliding behavior and the molecular level changes in gene expression associated with EMT. We focused on the downregulation of E-cadherin expression for two reasons. First, downregulation of E-cadherin is a well-established characteristic of cells undergoing EMT. In addition, we previously showed that downregulating E-cadherin using shRNA constructs enhances sliding and reduces CFD in MCF-10A cells; meanwhile, overexpressing E-cadherin in metastatic 231 cells inhibits sliding and increases CFD (17). Therefore, we hypothesized that the progressive enhancement in cell sliding during EMT may correspond with concomitant decreases in E-cadherin expression. To test this hypothesis, we quantified the TGFβ-induced changes in E-cadherin expression by Western blot (Fig. 3A). The results confirm that E-cadherin expression decreases gradually over time following TGFβ treatment. Both doses of TGFβ triggered downregulation of E-cadherin, with the higher dose of TGFβ having a quicker effect on reducing E-cadherin expression (Fig. 3B).

**Figure 3.**
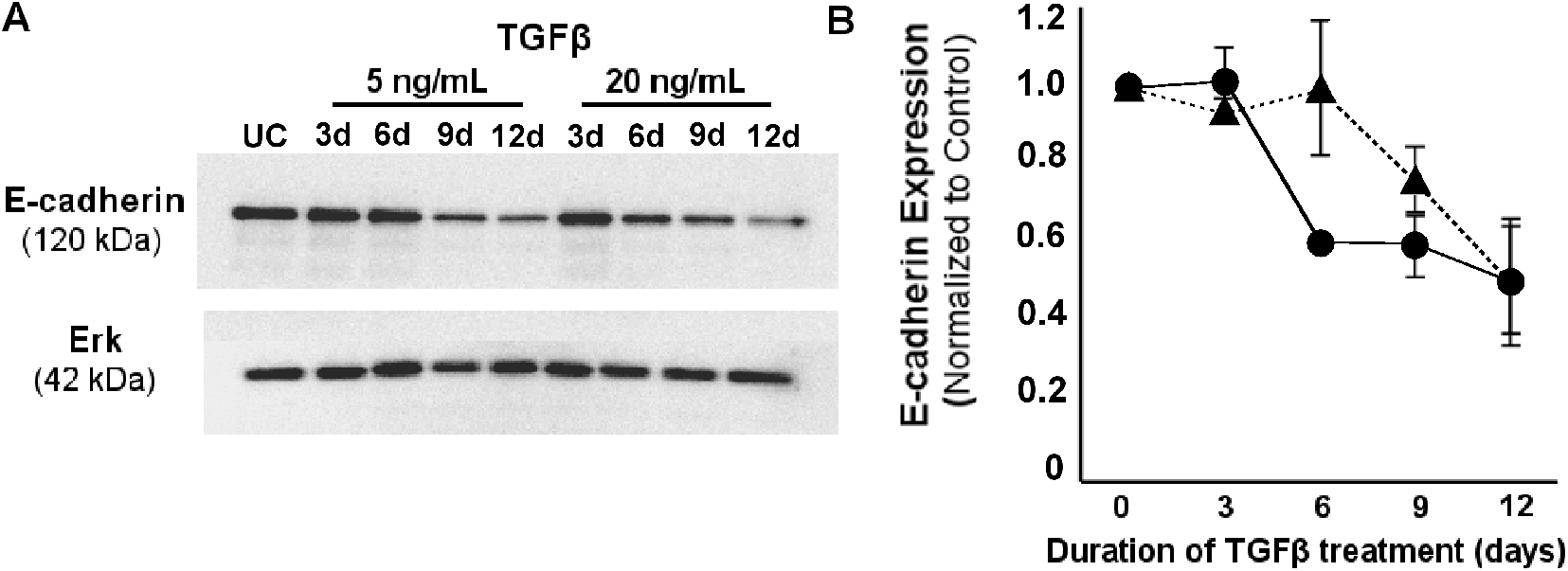
Extent of TGFβ-induced EMT. (*A*) E-cadherin expression was measured by Western Blot in TGFβ-treated and untreated (control) cells. (*B*) Quantification of the ratio of E-cadherin:Erk expression for cells treated with 5 ng/ml (triangle) or 20 ng/ml (circle) TGFβ. Values are reported relative to the E-cadherin:Erk expression ratio for untreated (UC) cells. Error bars denote S.E.M.

We probed the quantitative relationship between E-cadherin expression and cell sliding behavior across the different TGFβ treatments and the untreated control. Cells treated with 5 ng/mL TGFβ for 3 and 6 days and 20 ng/mL TGFβ for 3 days have approximately the same level of E-cadherin expression as untreated controls (Fig. 4). Yet, the CFD and sliding behavior are significantly different among these conditions. We conclude that in this regime of EMT, molecular changes independent of E-cadherin are mechanistically involved in enhancing cell-sliding behavior. In contrast, in the second regime of EMT where 20 ng/ml TGFβ is applied for 12 d, E-cadherin expression decreases by ~;50% and corresponds to a reduction in CFD and enhancement in cell sliding. In this portion of the EMT process, the loss of E-cadherin expression is correlated with an increase in the ability of cells to undergo contact-initiated sliding. Taken together, these results help to identify segments of the complex EMT process in which cell sliding behavior is enhanced independent of E-cadherin regulation and other segments in which downregulation of E-cadherin may facilitate cell sliding.

**Figure 4:**
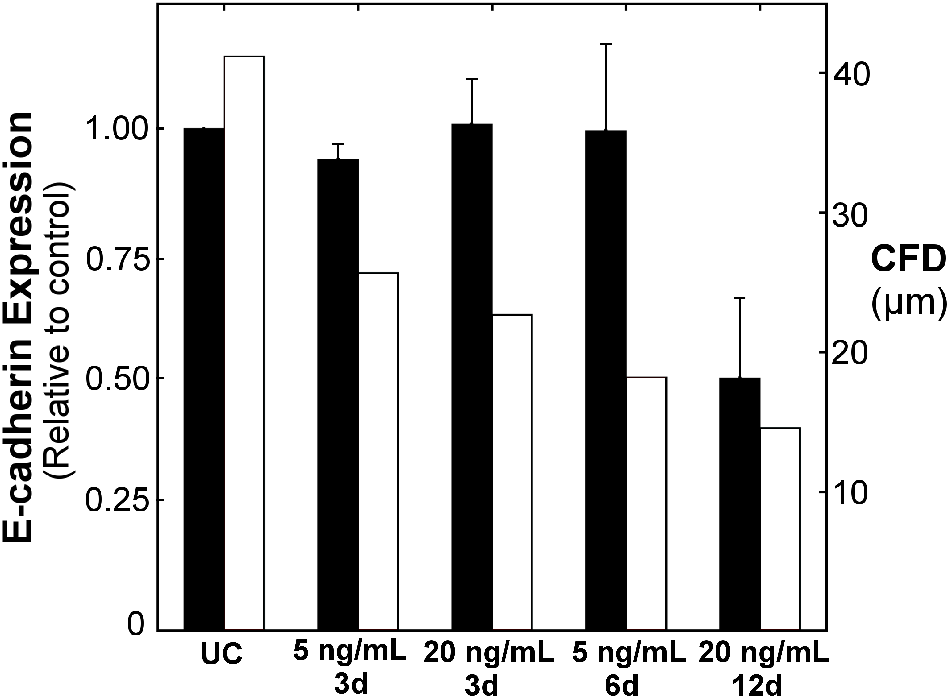
TGFβ-induced EMT promotes two regimes of invasive behavior. E-cadherin expression (black bars, left y-axis) and CFD values (white bars, right y-axis) are plotted for cells left untreated (UC) or treated with different doses and durations of TGFβ (x-axis).

## Conclusions

In summary, the results reported here provide three main insights into the quantitative relationship between EMT and invasive cell sliding behavior. First, EMT is a strong driver of the invasive cell sliding phenotype. When cells have progressed into EMT under maximal stimulation (20 ng/mL TGFβ for 12 days), the CFD drops to 15 µm. For comparison, we previously reported a CFD of 10 µm for triple-negative, claudin-low BT549 cells(17). Thus, exposing an otherwise non-invasive epithelial MCF-10A cell to 20 ng/mL of TGFβ for 12 days is sufficient to induce an invasive phenotype on par with highly metastatic breast cancer cells. Second, EMT enables cells to slide under extremely tight spatial constraints. The CFD of 15 µm under maximal TGFβ treatment is 2.7-fold narrower than the CFD of 41 µm for untreated control cells. To put this physical restriction in perspective, a CFD of 15 µm is comparable to the diameter of a single MCF-10A cell. Finally, partial progression through EMT leads to intermediate levels of invasive cell sliding behavior. Sub-maximal treatment with TGFβ using reduced dose and/or shorter duration of exposure promotes partial changes in cell morphology while concomitantly reducing CFD. The shift in CFD from 41 µm for untreated cells to 15 µm under maximal TGFβ stimulation is not a switch-like transformation but rather occurs through progressive shifts in sliding ability under ever tighter spatial constraints. These findings suggest a model in which fiber maturation and partial EMT work synergistically to promote invasiveness during cancer progression.

